# Paw placement during walking is altered by analgesic doses of opioids and post-surgical injury in mice

**DOI:** 10.1101/2021.12.15.472818

**Authors:** Victoria E. Brings, Maria A. Payne, Robert W. Gereau

## Abstract

Hind paw-directed assays are commonly used to study the analgesic effects of opioids in mice. However, opioid-induced hyper-locomotion can obscure results of such assays. We aimed to overcome this potential confound by using gait analysis to observe hind paw usage during walking in mice. We measured changes in paw print area following induction of post-surgical pain (using the paw incision model) and treatment with oxycodone. Paw incision surgery reduced the paw print area of the injured hind paw as the mice avoided placing the incised section of the paw on the floor. Surprisingly, oxycodone caused a tiptoe-like gait in mice, resulting in a reduced paw print area in both hind paws. Further investigation of this opioid-induced phenotype revealed that analgesic doses of oxycodone or morphine dose-dependently reduced hind paw print area in uninjured mice. The gait changes were not dependent on opioid-induced increases in locomotor activity; speed and paw print area had no correlation in opioid-treated mice, and other analgesic compounds that alter locomotor activity did not affect paw print area. Unfortunately, the opioid-induced “tiptoe” gait phenotype prevented gait analysis from being a viable metric for demonstrating opioid analgesia in injured mice. However, this work reveals an important, previously uncharacterized effect of treatment with analgesic doses of opioids on paw placement. Our characterization of how opioids affect gait has important implications for the use of mice to study opioid pharmacology and suggests that scientists should use caution when using hind paw-directed nociceptive assays to test opioid analgesia in mice.

## INTRODUCTION

Mice are widely used to study the behavioral effects of opioids, including analgesia, dependence, and respiratory depression [8; 13; 28; 31]. Antinociceptive properties of mu opioid agonists are often tested in mice using reflexive hot plate or tail flick assays [13; 31; 35]. To study opioid analgesia in tissue or nerve injury models in mice, injury-induced sensitization and its reversal with opioids are assessed by measuring sensitivity to stimuli directed to the affected body region, which is often the hind paw [12]. Most commonly used hind paw-directed assays entail the quantification of a withdrawal response to the application of noxious stimuli (heat, cold, touch) to the hind paw [5; 12]. These assays reflect evoked responses to an exogenous stimulus and do not reflect naturalistic pain behaviors [12; 32]. They also measure reflexive withdrawal responses, which may not accurately represent pain or the effects of analgesics [15; 18]. In addition to having potentially flawed face validity, hind paw-directed evoked pain assays are particularly challenging to perform on opioid-treated mice due to increases in locomotor activity induced by mu opioid agonists [10; 34; 35].

Because of the shortcomings of evoked withdrawal tests, alternative approaches are needed to assess opioid analgesia in mice. We and others have used a variety of non-reflexive behavioral assays to assess pain sensitivity and analgesia in mice, such as voluntary wheel running, home cage lid hanging, and gait analysis [21; 30; 32; 37; 41]. To achieve our goal of measuring behavior affected by injury and reversed by opioids in mice, in the present study we use the CatWalk gait analysis system to observe hind paw usage during voluntary walking behavior [1; 21; 30; 37]. Gait analysis provides a means of assessing hind paw usage with a non-reflexive behavioral assay that would be technically feasible with a hyperactive mouse.

Using the CatWalk system, we measured changes in paw print area following induction of post-surgical pain with the paw incision injury model and treatment with oxycodone in mice. We hypothesized that paw incision injury would produce alterations in paw placement during walking that could be reversed with oxycodone. We found that paw incision injury reduces the print area of the injured paw. Surprisingly, however, analgesic doses of oxycodone induce a “tiptoe” phenotype that reduces the print area of all paws. We further characterize how oxycodone and morphine dose-dependently alter paw placement during walking in mice. The reduction in paw print area is independent of walking speed and is specific to opioids, as mice treated with other analgesics do not show altered paw print area. Overall, this work demonstrates the effect of opioid treatment on paw usage and has implications for the use of hind paw-directed sensory assays in opioid-treated mice.

## MATERIALS AND METHODS

### Animals

All procedures used in this study were approved by the Institutional Animal Care and Use Committee at Washington University in St. Louis School of Medicine. C57BL/6 mice bred in-house were maintained in a facility with a 12-h light:dark cycle (6 AM-6 PM lights on) and given food and water *ad libitum*. Animals were group-housed with 3 to 5 mice per cage. Mice were reared on 1/4-inch corn cob bedding from birth until transfer to a separate room at least 3 days prior to testing, where they were then house with 1/8-inch corn cob bedding. Behavior was tested during the light phase at 8 to 12 weeks of age. Male and female mice were always tested separately. Animals were randomly assigned to each condition. Different test groups (injury, drug treatment) had equal average weights. Cages had mixes of animals from different test groups. Female mice were used to test the effects of paw incision, oxycodone, morphine, THC, and fenobam; male mice were tested for their responses to oxycodone in the CatWalk experiments. Experimenters were blinded to drug treatment or surgical condition until after data were analyzed.

### Drugs

Oxycodone (USP, Rockville, MD) was dissolved in a vehicle of saline (0.9% NaCl; Hospira, Lake Forest, IL), and morphine (10 mg/mL; Hikma, Eatontown, NJ) was diluted in saline; each was delivered s.c. in a volume of 5 mL/kg body weight 30 min before testing behavior. Δ^9^-tetrahydrocannabinol (THC; 200 mg/mL; National Institute on Drug Abuse Drug Supply Program, Bethesda, MD) was diluted with 95% ethanol followed by a mix of Kolliphor EL (Sigma-Aldrich, St. Louis, MO) and saline to achieve a final vehicle of 5% 95% ethanol, 5% Kolliphor EL, and 90% 0.9% saline, delivered s.c. in a volume of 5 mL/kg body weight 30 min before testing behavior. Fenobam (SCYNEXIS, Durham, NC) was dissolved in a vehicle of 100% DMSO (Sigma-Aldrich), delivered i.p. in a volume of 20 μL 5 min before testing behavior. For all drugs, testing was done within 30 min (30-60 min post-delivery of oxycodone, morphine, Δ^9^-THC; 5-35 min post-delivery of fenobam). Doses were chosen based on previous reports that analgesia and/or antinociception can be achieved at the indicated doses for oxycodone [2; 14; 25; 29; 38; 39], morphine [25; 29], fenobam [24; 26; 27], and Δ^9^-THC [3; 9; 22]. All injections were made with a 0.3 mL insulin syringe with a 1/2 inch 29 gauge needle (Exelint, Redondo Beach, CA). Mice were returned to home cage following injection and were kept there until testing.

### Paw incision surgery

Paw incision surgery was performed on the right hind paw (**Figure 1A**), following previous published procedures [11; 40]. While under isoflurane anesthesia, the surface of the right hind paw was sterilized and a 5-mm incision was made through the skin starting 2 mm from the heel using a type 11 scalpel blade (McKesson, Richmond, VA). The flexor digitorum brevis muscle was pulled up to isolate, with rough movement beneath the muscle to promote sensitization, and an approximately 3-mm vertical incision was made through the muscle without severing the muscle. The skin wound was closed with two 6-0 silk sutures with a tapered needle (Ethicon, Guaynabo, Puerto, Rico), and antibiotic ointment (McKesson) was applied. The sham surgery was identical, except no incisions were made; mice were anesthetized, skin was sterilized and sutured, and ointment was applied. Sham and incision animals were housed in mixed cages.

**Figure 1.**
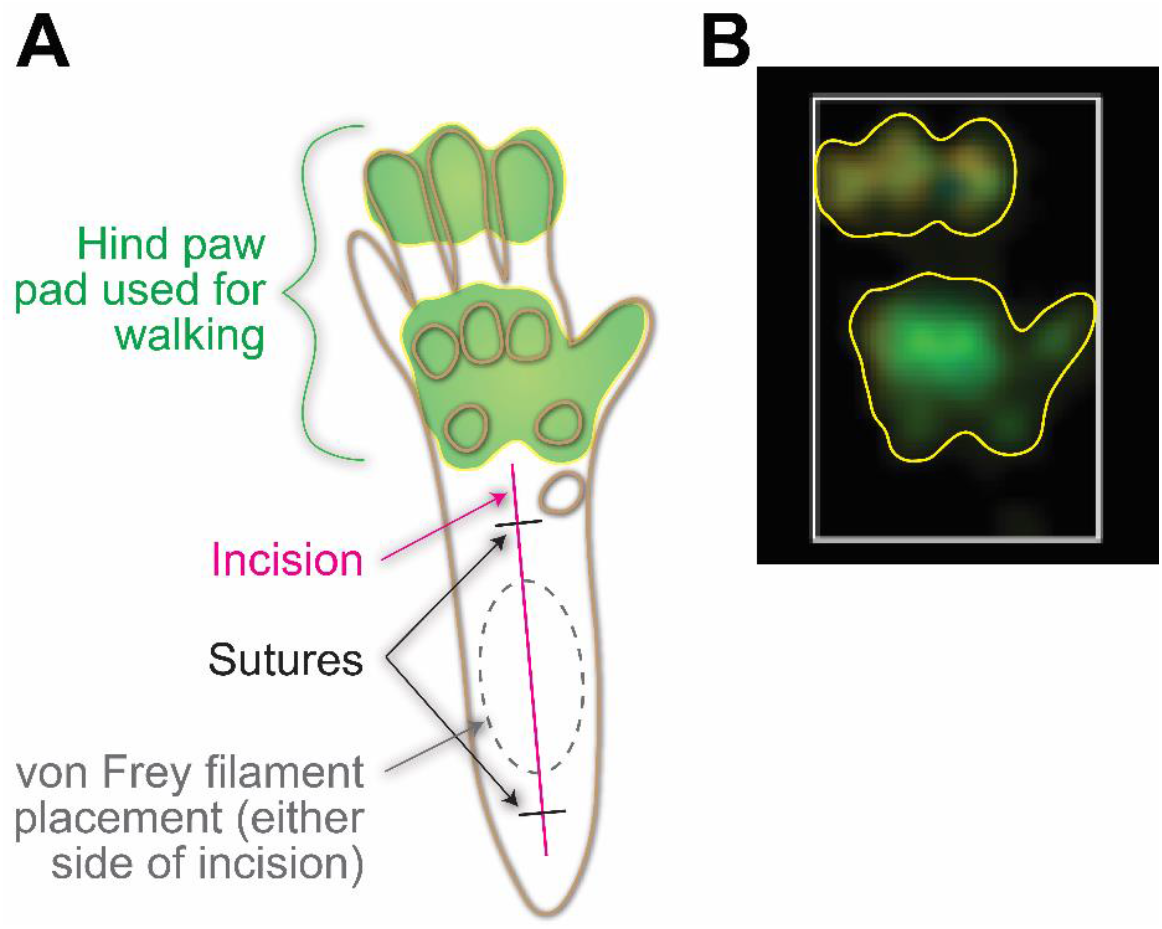
Diagram of hind paw showing locations used for sensory testing and incision injury. (A) An outline of a mouse hind paw showing circles that correspond to raised sections of the foot pad (brown). The approximate size and location of the incision (magenta) and the two sutures that close the wound (black) are also shown, along with the area that the von Frey filament was placed during sensory testing (gray dashed circle). The area of the hind paw that makes contact with the ground is indicated (green). (B) A paw print image produced by the Noldus CatWalk software (white box surrounding paw print added by software). Yellow outlines added to (A) and (B) indicate the outline of the paw print captured by the software and the corresponding hind paw anatomy that generated the print.

### Electronic von Frey

Mechanical allodynia was measured using an electronic von Frey anesthesiometer (IITC Life Science, Woodland Hills, CA) [33]. Mice were placed on a stainless steel mesh platform elevated approximately 30 cm from the bench top. Mice were isolated in individual clear acrylic containers (10 cm x 10 cm x 15 cm high). Black plastic dividers between the containers prevented mice from seeing each other. After at least one day of acclimation to the apparatus, on test day mice were allowed to habituate for 1 hour prior to testing, with the experimenter in the room for the 30 min prior to testing. Alternating hind paws, a total of 5 measurements were taken per paw, with at least 1 min between measurements for a single paw. All measurements were taken for a single animal within a 15 min period. On the day of testing post-surgery, approximately 15 min after acquiring baseline data the animals were then injected with the test compound and tested again for post-drug responses 30 min later. The blunt tip of the anesthesiometer was applied to the hind paw approximately 4-5 mm from the heel (which corresponds to the space between the two sutures on the hind paw in sham and injured animals, **Figure 1A**), and the amount of force applied before the animal withdrew their hind paw was recorded. Of the 5 measurements taken per paw, the highest and lowest measurements were excluded, and the remaining 3 measurements were averaged.

### CatWalk gait analysis

The Noldus CatWalk XT gait analysis system (Noldus Information Technology, Leesburg, VA) was used to analyze the paw prints of mice [1; 21; 30; 32; 37]. In a dark room, mice walked across a glass platform elevated 118 cm from the ground with a camera positioned 32 cm underneath the glass, facing the underside of the animal. Mice walked across the 6 cm wide pathway and light passing perpendicularly through the glass was refracted when weight was placed on the glass, illuminating the section of the paws that apply that weight (**Figure 1A**). Paw prints were imaged by the camera and measured using the Noldus CatWalk XT 10 software (**Figure 1B**). The paw print area refers to the total area of the glass in contact with the paw; the space between toes is not included in the area measurement. The paw print area was measured at the point in the step when the area of contact with the glass is maximal for that paw print. Analgesic studies in incision animals were performed in mice that had a relative paw print area in the ipsilateral paw that was no greater than 80% than of the contralateral paw.

### Imaging mouse walking behavior

Mice were placed on a glass platform elevated 21 cm from the bench top and allowed to walk along a 10 cm wide pathway while they were imaged with a Google Pixel 4 (Google, Mountain View, CA).

### Statistics

Data were analyzed using GraphPad Prism 9.3.0 (GraphPad Software, San Diego, CA). Data were analyzed with one- or two-way ANOVA tests, followed by Dunnett’s or Tukey’s multiple comparisons tests, respectively. When two groups were being compared, t-tests were used (paired or unpaired, as indicated below). A simple linear regression was used to test the correlation between paw print area and speed. Statistical values and tests used are described in the test below. Significant main effects are reported in the text and significant post-hoc test results are indicated in the figures.

## RESULTS

### Oxycodone reverses incision-induced mechanical allodynia but not reduced paw print area of injured paw

Consistent with prior reports, we found that paw incision produced robust sensitization to mechanical stimuli in mice as assessed using von Frey filaments one day after injury (**Figure 2A**). After treatment with oxycodone (10 mg/kg s.c.) on day 1 post-surgery, the withdrawal thresholds were no longer significantly different from pre-injury baseline values (F_6, 60_ = 7.097, p < 0.0001, two-way ANOVA interaction effect).

**Figure 2.**
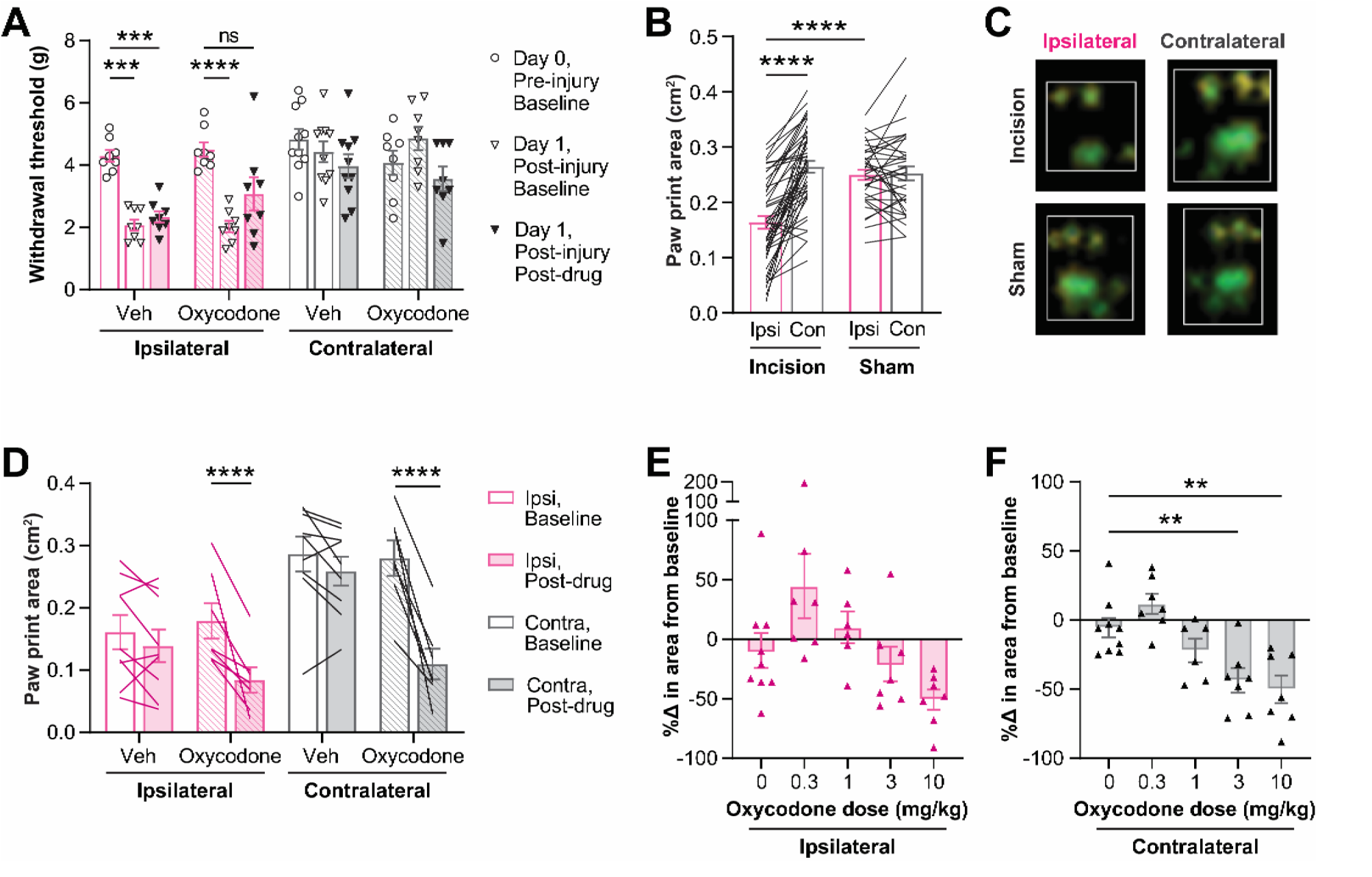
Oxycodone reverses mechanical allodynia but not paw print changes following paw incision surgery. A) Withdrawal threshold was measured in contralateral and ipsilateral hind paws before paw incision surgery, 1 day after surgery, and 30 min after treatment with 10 mg/kg of oxycodone or vehicle on day 1 after surgery. B) Paw incision surgery reduced the paw print area of the injured (ipsilateral) hind paw, relative to the animals’ contralateral paw or to sham animals’ ipsilateral paws. C) View of the hind paw print image collected by the CatWalk system when the paw makes maximum contact with the glass in a paw incision injured mouse. D) The paw print area was measured in both hind paws twice on the day after surgery: before (baseline) and after (post-drug) treatment with 10 mg/kg of oxycodone or vehicle. E-F) On day 1 after paw incision surgery, the percent change in paw print area from baseline to post-drug (30 min after treatment with indicated oxycodone doses) in the ipsilateral (E) and contralateral (F) hind paws. All oxycodone administration was s.c., 30 min prior to post-drug test. Mean ± SEM. Showing results of post-tests following 2-way ANOVAs for A (p<0.0001), D (p<0.0001), and F (p<0.0001); p< ^**^0.01, ^***^0.001, ^****^p<0.0001. B) Unpaired t-test of sham vs. incision ipsilateral; paired t-test of incision contralateral vs. ipsilateral; ^****^p<0.0001.

Paw incision alters paw placement by inducing paw guarding behavior in rats and mice [4; 40]. Therefore, we hypothesized that the CatWalk gait analysis system, which evaluates paw placement during walking, would detect altered paw usage in injured mice. We performed paw incision surgery and analyzed walking using the CatWalk system one day post-surgery. Paw print area was measured for each hind paw at the point of the step when the paw makes maximum contact with the glass floor. Paw incision surgery reduced the paw print area of the injured (ipsilateral) hind paw, relative to the animals’ contralateral paw (p < 0.0001, paired t-test) or to sham animals’ ipsilateral paws (p < 0.0001, unpaired t-test; **Figure 2B**). Viewing images of the paw prints shows that the reduction in paw print area is due to an avoidance of placing weight on the proximal section of the paw, closer site of the incision (**Figure 2C**).

Next, we assessed if oxycodone could reverse the incision-induced reduction in paw print area (**Figure 2D**). As in **Figure 2B**, paw print area was reduced in injured hind paws relative to contralateral paws one day post-surgery. We hypothesized that oxycodone would increase the contact area of the injured paws. Surprisingly, however, oxycodone (10 mg/kg s.c.) decreased the paw print area of both uninjured and injured hind paws (F_4, 31_ = 10.64, p < 0.0001, two-way ANOVA interaction effect). In other injury models, different doses of oxycodone were required to reverse sensitization phenotypes measured with different behavior tests [25; 39]. Therefore, we sought to determine if a lower dose of oxycodone would reverse the impaired paw placement in the incised paws without affecting the contralateral paws (**Figure 2E-F**). No dose of oxycodone significantly altered paw print area of the injured paw relative to vehicle (i.e. 0 mg/kg of oxycodone) treatment (**Figure 2E**); however, 3 and 10 mg/kg of oxycodone significantly decreased the area of the contralateral paw prints relative to vehicle (F_4, 31_ = 9.589, p < 0.0001, one-way ANOVA; **Figure 2F**).

### Opioids dose-dependently decrease paw print area during walking

Based on the finding of oxycodone’s effect on contralateral paw print area, we decided to further investigate the effect of oxycodone on paw placement in uninjured mice. Using a range of analgesic doses of oxycodone [2; 14; 25; 29; 38; 39], we assessed paw placement during walking before and after oxycodone treatment, as in **Figure 2D**. We found that oxycodone caused dose-dependent reductions in the footprint area of the hind paws in both female (F_4, 45_ = 12.48, p < 0.0001, two-way ANOVA interaction effect; **Figure 3A**) and male (F_4, 40_ = 19.86, p < 0.0001, two-way ANOVA interaction effect; **Figure 3B**) mice, and the effect of oxycodone was greater than that of vehicle in females (F_4, 45_ = 11.67, p < 0.0001, one-way ANOVA; **Figure 3C**) and males (F_4, 40_ = 26.65, p < 0.0001, one-way ANOVA; **Figure 3D**), as well. In females the hind paw print area was reduced after 3 or 10 mg/kg oxycodone treatment (**Figure 3A**), and in males the hind paw print area was reduced after 1, 3, or 10 mg/kg oxycodone treatment (**Figure 3B**), relative to their baseline print areas. Front paw print area was also significantly reduced by oxycodone in a dose-dependent fashion in both females (F_4, 45_ = 11.27, p < 0.0001, two-way ANOVA interaction effect; **Figure 3E**) and males (F_4, 40_ = 7.980, p < 0.0001, two-way ANOVA interaction effect; **Figure 3F**). Significant reductions relative to baseline were seen after 3 and 10 mg/kg oxycodone treatment in both female and male front paw print area, and, as with the hind paws, the change in paw print area induced by oxycodone was significantly greater than that induced by vehicle in females (F_4, 45_ = 7.771, p < 0.0001, one-way ANOVA; **Figure 3G**) and males (F_4, 40_ = 7.539, p = 0.0001, one-way ANOVA; **Figure 3H**).

**Figure 3.**
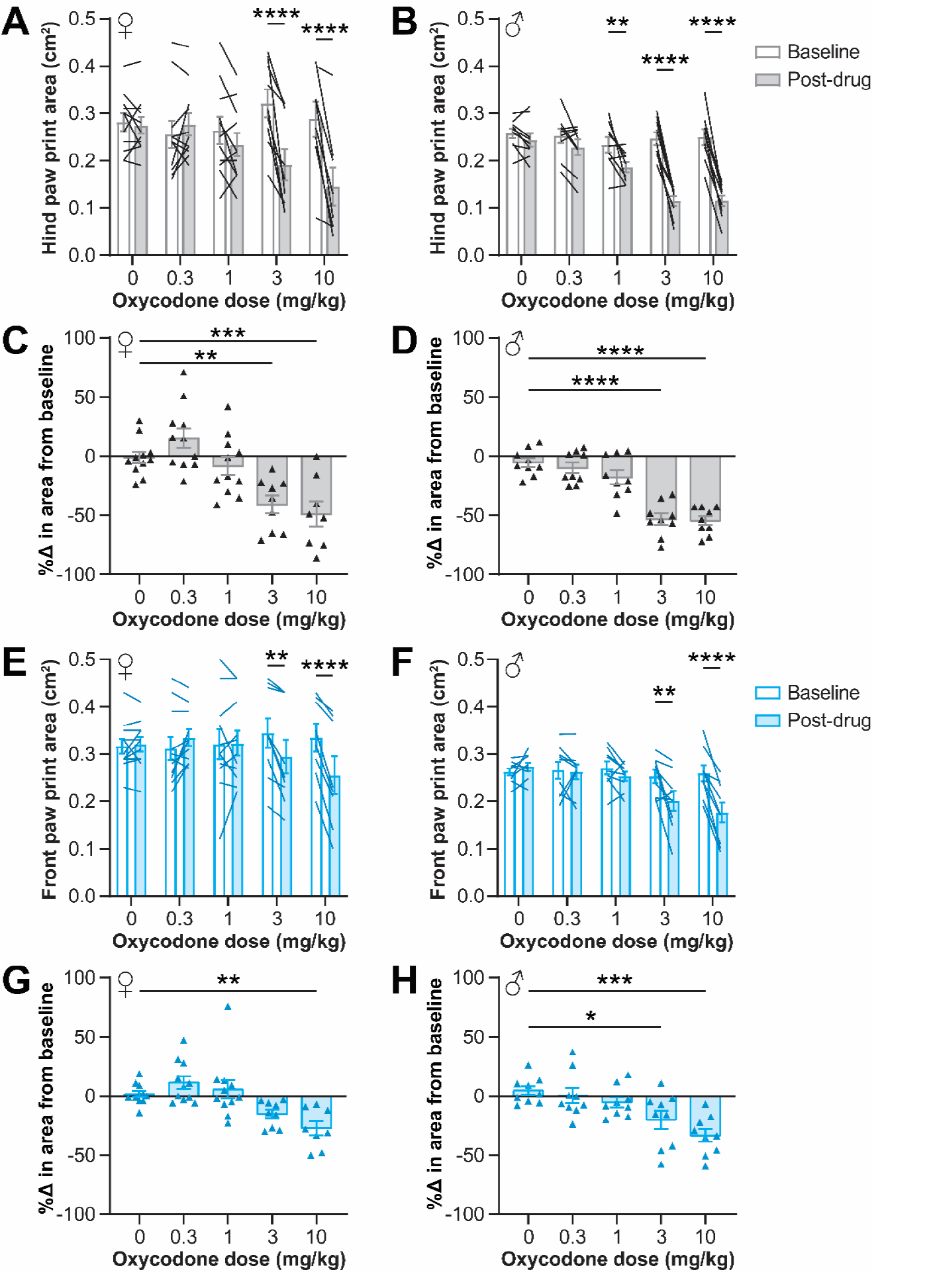
Analgesic doses of oxycodone reduce paw print area in female and male mice. Analgesic doses of oxycodone reduced hind paw (A-D) and front paw (E-H) print area in uninjured female (A, C, E, G) and male (B, D, F, H) mice. (A-B,E-F) The area of the paws at the point of maximum paw contact before (baseline) and after (post-drug) oxycodone administration. Graph legend on (B) applies to (A) and (B); graph legend on (F) applies to (E) and (F). (C-D,G-H) The percent change between baseline and post-drug for each animal. Mean ± SEM. Showing results of post-tests following 2-way ANOVAs for A (p<0.0001), B (p<0.0001), E (p<0.0001), and F (p<0.0001) and 1-way ANOVAs for C (p<0.0001), D (p<0.0001), G (p<0.0001), and H (p=0.0001); p< ^*^0.05, ^**^0.01, ^***^p<0.001, ^****^p<0.0001.

To assess how oxycodone alters paw placement to reduce paw print area, we examined the paw print images and images of animals walking following oxycodone treatment. Images of the paw prints when the maximum contact is made with the glass showed that mice favored the toes and balls of their feet after treatment with 10 mg/kg of oxycodone relative to their walking behavior at baseline (**Figure 4A**). While observing mice at the moment when the hind paw first lands during a step, we saw that vehicle-treated mice (**Figure 4B**) placed their full hind paw pad down on the glass and that 10 mg/kg oxycodone-treated mice (**Figure 4C**) avoided placing the more proximal section of their paws (i.e. their heels) down on the glass. Oxycodone-treated mice also showed rigid, vertically pointed tails and, by adopting a tiptoe-like posture while walking, were more elevated from the platform. The avoidance of heel placement seen following oxycodone treatment could thus produce a reduction in paw print area.

**Figure 4.**
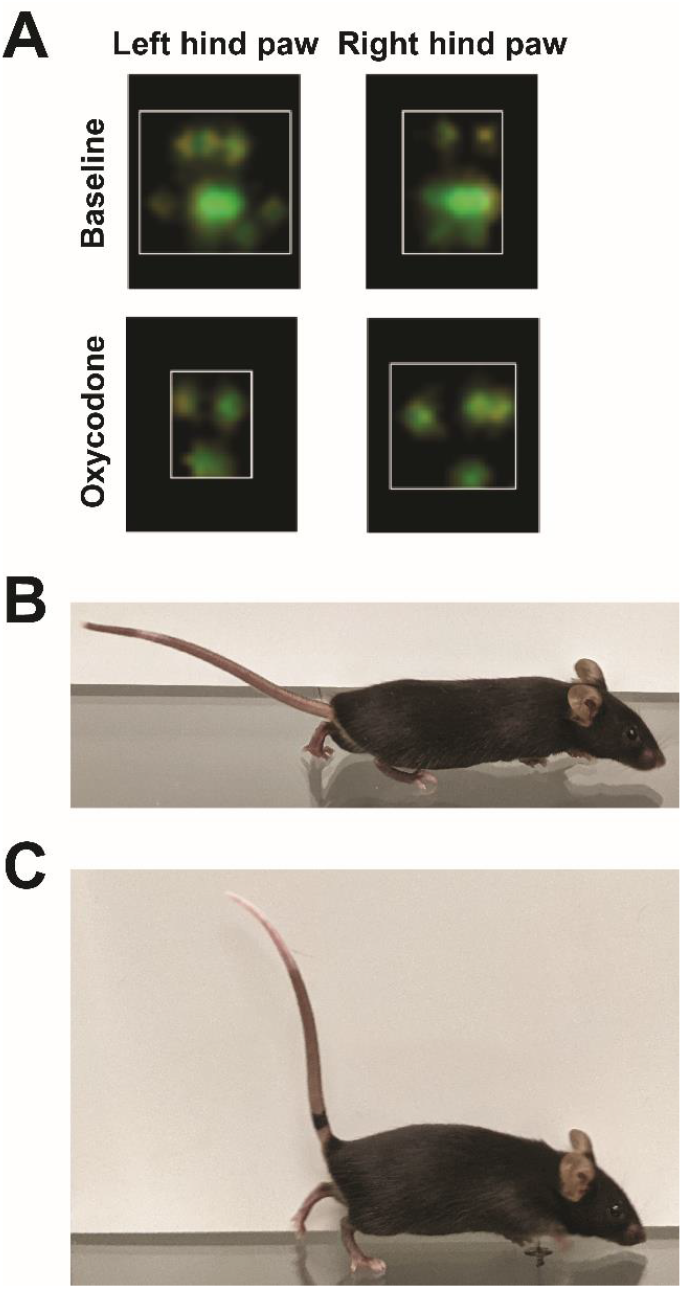
Analgesic doses of oxycodone reduce paw print area by inducing tiptoe walking. (A) Paw prints were imaged by the Noldus CatWalk software. White boxes show edges of the paw print. Mice were photographed while walking on an elevated glass platform 30 min after treatment with vehicle (B) or 10 mg/kg of oxycodone (C). The oxycodone-treated mouse shows incomplete placement of the hind paw on the glass, elevated body position relative to the platform, and a rigid Straub tail.

To determine if the effect on paw placement found with oxycodone could be see with another analgesic mu opioid receptor agonist, we examined paw placement on the CatWalk before and after morphine treatment. Morphine also dose-dependently decreased paw print area. As with oxycodone, 10 mg/kg of morphine (s.c.) reduced paw print area relative to baseline (F_4, 28_ = 2.840, p = 0.0428, two-way ANOVA effect of drug; **Figure 5A-B**), but the effect sizes were smaller than at the same doses of oxycodone (**Figure 3A,C**).

**Figure 5.**
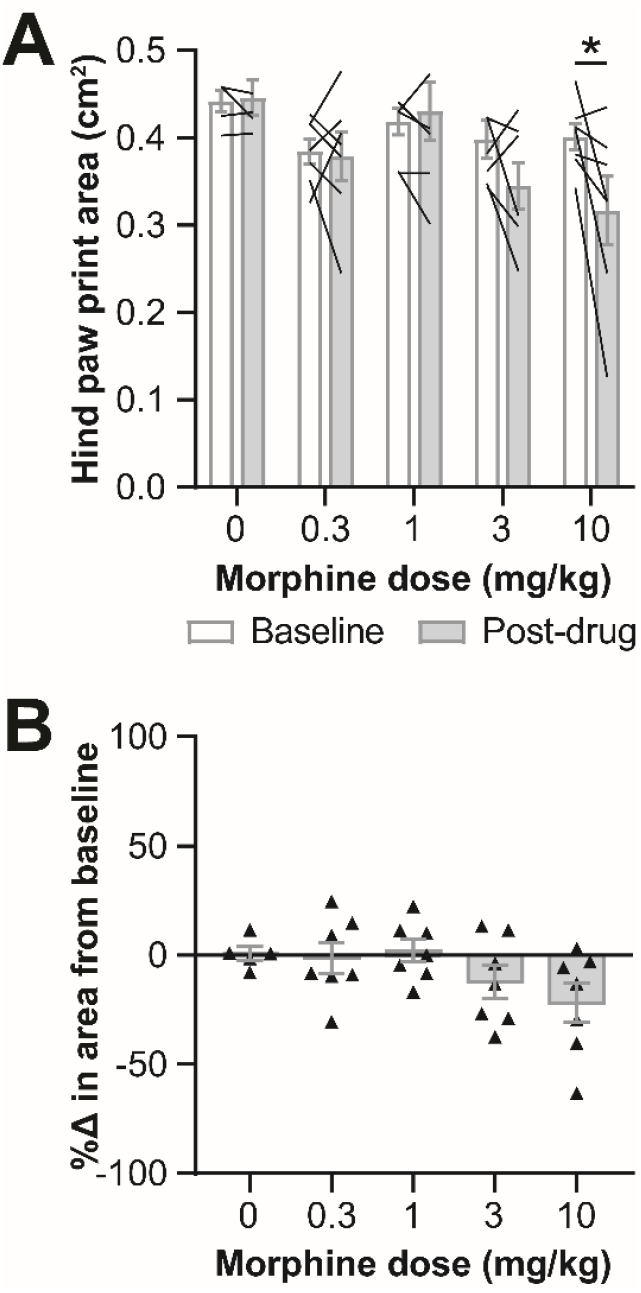
Analgesic doses of morphine reduce hind paw print area during walking in female mice. (A) The area of the hind paws at the point of maximum paw contact before (baseline) and after (post-drug) morphine administration (s.c., 30 min before test). (B) The percent change in paw print area between baseline and post-drug for each animal. Mean ± SEM. Showing results of post-tests following a 2-way ANOVA for A (p=0.0428); p< ^*^0.05.

### Altered walking speed does not drive reduced paw print area

The oxycodone doses used here significantly increase distance covered in an open field [10] and, thus, with time being a constant, average speed in an open field is increased. Therefore, we asked whether altered walking speed could cause the opioid-induced tiptoe-like gait. We found that the altered paw placement was not simply a product of increased locomotion, as we saw no correlation between the paw print area and the speed of the mice (**Figure 6A-B**).

**Figure 6.**
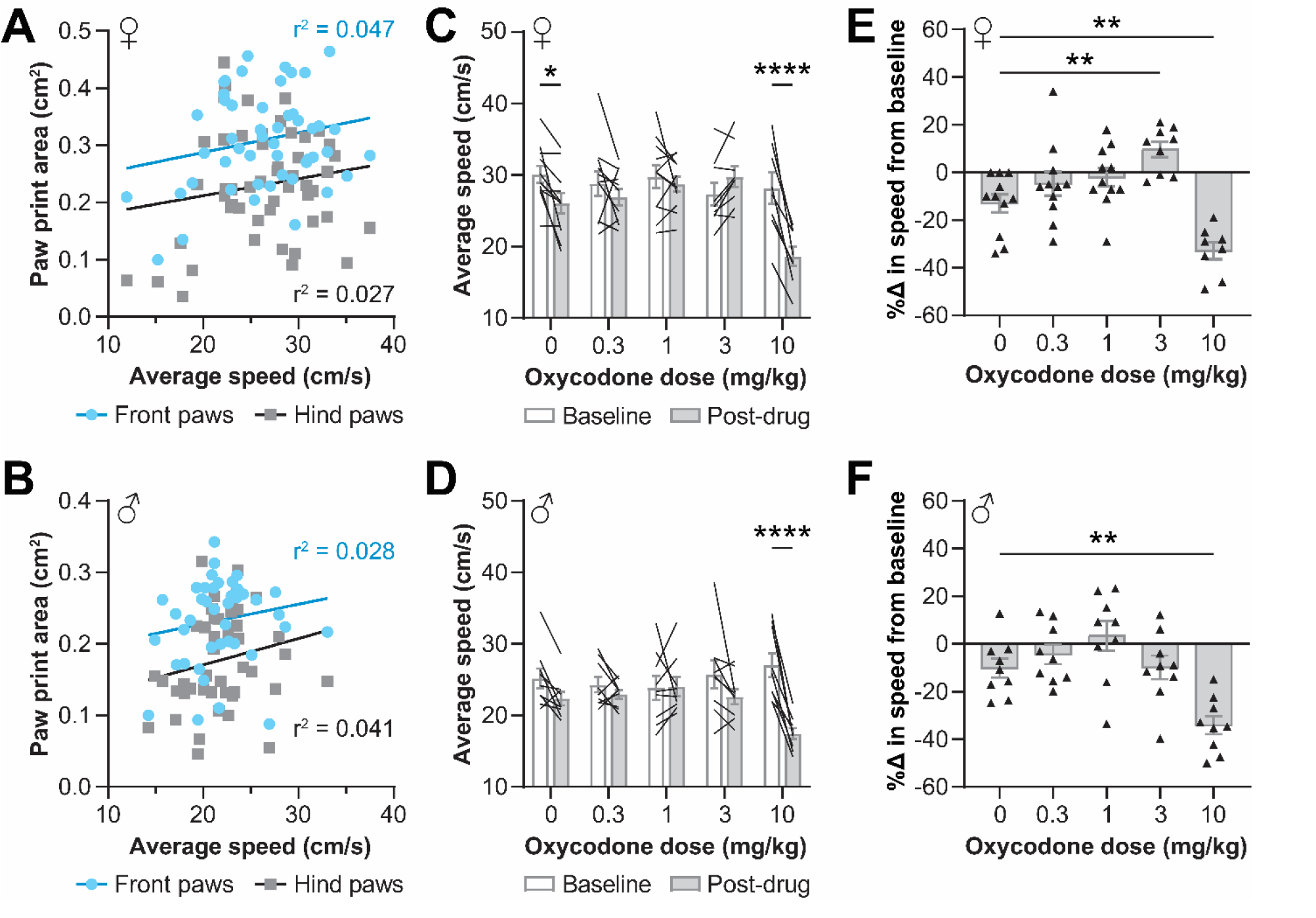
Oxycodone’s effect on walking speed does not drive altered paw print area. Speed and front or hind paw print area had no correlation in oxycodone-treated female (A) or male (B) mice. The average speed during walking on the CatWalk platform before (baseline) and after (post-drug) oxycodone administration in females (C) and males (D). The percent change between baseline and post-drug for each female (E) and male (F) mouse. Showing results of linear regression for (A-B) and post-tests following 1-way ANOVAs for C (p<0.0001), D (p=0.0003), E (p<0.0001), and F (p<0.0001); p< ^*^0.05, ^**^0.01, ^****^p<0.0001. (C-F) Mean ± SEM.

Nearly all vehicle-treated mice had slower average speeds on the CatWalk during their post-drug trials relative to their baseline trials in female mice (F_4, 45_ = 9.736, p < 0.0001, two-way ANOVA interaction effect; **Figure 6C**), and both female and male mice (F_4, 40_ = 6.671, p = 0.0003, two-way ANOVA interaction effect; **Figure 6D**) were significantly slower during their post-10 mg/kg of oxycodone trials relative to baseline, as well. Notably, the reduction in speed during post-drug trials was significantly greater in the mice treated with 10 mg/kg of oxycodone relative to the change in speed in mice treated with the vehicle in both females (F_4, 45_ = 12.88, p < 0.0001, one-way ANOVA; **Figure 6E**) and males (F_4, 40_ = 8.954, p < 0.0001, one-way ANOVA; **Figure 6F**). Interestingly, in female mice 3 mg/kg of oxycodone increased walking speed relative to baseline (**Figure 6E**). Hind paw print area was decreased at both 3 and 10 mg/kg of oxycodone in females (**Figure 3A,C**), but speed was increased at 3 mg/kg and decreased at 10 mg/kg of oxycodone. Additionally, the effects of oxycodone on hind paw area in males (**Figure 3B,D**), particularly the equivalent effects of 3 and 10 mg/kg on paw placement, did not match the effects of oxycodone on walking speed (**Figure 6F**), as only 10 mg/kg of oxycodone affected walking speed in male mice. The reduction in walking speed induced by 10 mg/kg of oxycodone seen in both sexes is notable, as it suggests the increase in distance traveled in an open field at this dose may be due to an increase in time spent walking rather than an increase in walking speed. Together, these data support the conclusion that the change in speed caused by oxycodone treatment did not drive the change in paw placement.

As an additional exploration into the effect of walking speed on paw placement, we also tested the effects of two other non-opioid analgesic drugs on paw print area in female mice. We chose to test analgesic compounds that differentially affect open field activity. Fenobam, an mGlu5 receptor negative allosteric modulator, reduces pain behaviors and increases open field activity [26]. Δ^9^-THC, a cannabinoid type 1 and type 2 receptor agonist, provides anti-nociception and decreases open field activity [3; 9; 22]. Neither fenobam (30 mg/kg, i.p.) nor Δ^9^-THC (3 mg/kg, s.c.) affected paw print area differently from their vehicle-treated counterparts (**Figure 7A-B**). Similarly to the oxycodone-treated mice, the effects on speed also did not match their respective effects on previously-reported open field activity. Fenobam and vehicle treatment had equivalent effects on speed (**Figure 7C**), and mice treated with Δ^9^-THC did not have the same reductions in walking speed as the vehicle-treated mice (p = 0.0192, unpaired t-test; **Figure 7D**).

**Figure 7.**
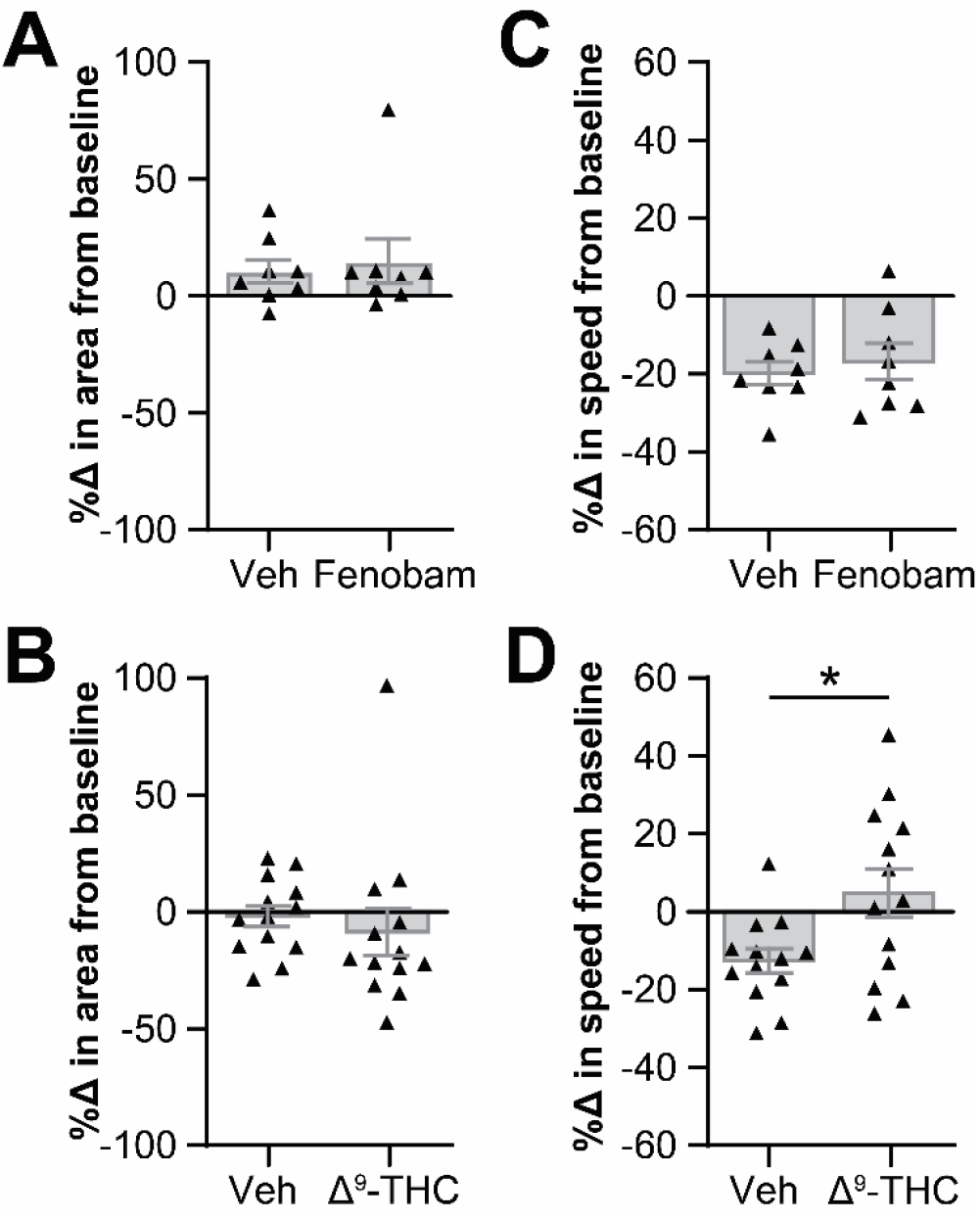
Non-opioid analgesics fenobam and Δ^9^-THC do not affect paw print area. The percent change in hind paw area (A-B) and average walking speed (C-D) between baseline and post-drug walking in mice treated with (A,C) fenobam (30 mg/kg, i.p., 5 min before test) and (B,D) Δ^9^-THC (3 mg/kg, s.c., 30 min before test). Mean ± SEM. Unpaired t-test in D; ^*^p<0.05.

## DISCUSSION

Overall we found that paw incision surgery reduced contact of the incised section of the paw during walking on the CatWalk platform, resulting in a decrease in the paw print area of the injured hind paw. Treatment with analgesic doses of opioids induced a tiptoe-like gait in injured and uninjured mice, causing reduced paw print area of both hind paws. Therefore, paw print area during walking cannot be used to as a phenotypic demonstration of post-injury sensitization that can be reversed by opioids in mice.

Within the results showing that opioids induce tiptoe walking, notable differences were seen between some subgroups. At the same doses, morphine had smaller effects on paw print area (**Figure 5A-B**) than oxycodone did (**Figure 3A,C**). A similar difference in the efficacy of equal doses has been noted for these two opioids in other phenotypes, e.g. morphine causes less severe respiratory depression than an equal dose of oxycodone in CD-1 mice [16]. Morphine has also been shown to be less analgesic in injured mice and has reduced anti-nociceptive efficacy in uninjured mice, relative to the same doses of oxycodone [25; 29]. Additionally, females and males responded differently to the same oxycodone doses. In males 1 mg/kg of oxycodone reduced hind paw print area relative to baseline (**Figure 3B**), whereas females were only affected at a dose as low as 3 mg/kg of oxycodone (**Figure 3A**). The sex differences in the effects of oxycodone on paw print area echo the body of literature demonstrating the enhanced sensitivity of male mice to the effects of opioids relative to female mice, including opioid-induced antinociception, tolerance, and locomotor activity [7; 10; 23].

In addition to the effects of opioids on paw print area, this work demonstrates an effect on walking speed that has not previously been quantified. Here, we show that male and female mice have a greater decrease in walking speed after treatment with 10 mg/kg of oxycodone than their vehicle-treated counterparts (**Figure 6C-F**). This dose of oxycodone was previously reported to increase total distance traveled in an open field in both sexes [10]. An increase in distance traveled is typically interpreted as an increase in average speed for the whole experiment, as the time of an open field test is kept constant. However, mice spend large portions of the open field period immobile. Our results suggest that the 10 mg/kg oxycodone-induced increase in distance covered may instead be driven by an increase in the percent of time spent mobile, with the walking speed (the average speed during mobile periods, not during the whole course of the test) actually being lower than that of vehicle-treated mice. An increase in the percent of time spent walking would also lead to more distance covered, a result that is often interpreted as being due to faster walking. As researchers explore the side effects of opioids in mice, including locomotor sensitization, our results on the effects of oxycodone on speed may help people better evaluate their data. Similar differential effects were shown here for THC and fenobam; the average walking speed in CatWalk did not align with the average speed previously shown in an open field, as calculated using the total distance covered. CatWalk and open field experiments have different conditions (e.g. floor material, lighting) that could contribute to different walking behaviors in each type of arena, but it would be interesting to parse out the effects on walking speed and time spent walking in an open field after treatment with these drugs as well.

One other mouse model that has shown tiptoe walking is the *twy* Yoshimura mouse, but the cause of tiptoe walking in this animal is a spinal injury arising from spinal ligament ossification and is unlikely to share a causal mechanism with acute opioid-induced tiptoe gait [17]. Others have anecdotally noted the presence of tiptoe-like walking induced by buprenorphine [19; 20], but none have quantified this behavior, and the cause is unknown.

One known side effect of mu opioid receptor agonist treatment that could cause altered paw placement is muscle rigidity. Rigid extension of the hind limbs resulting from subcutaneous morphine has been noted in mice [6], and was quantified in hind limb electromyographic activity studies using rats treated with subcutaneous alfentanil [36]. Muscle stiffness in the hind limbs could cause the back legs to be extended with minimal bending at the knee; with the legs stretched backward, the heels of the hind paw would not make contact with the walking surface, leading to a reduced hind paw print area. At higher oxycodone doses, this rigid extension may also occur in the fore limbs, leading to the reduced paw print area seen in the front paws at higher doses. Similarly, muscle rigidity is seen in the tail (commonly referred to as the Straub tail effect) in a dose-dependent fashion [6] that correlates well with the dose-dependent effect of tiptoe walking (see **Figure 4C** for an example of both phenotypes presented simultaneously), supporting the potential conclusion that muscle rigidity drives this effect.

An interesting effect on speed is seen in females treated with oxycodone; compared to their baseline speed, post-drug walking speed dose-dependently increases with oxycodone treatment up to 3 mg/kg, but is significantly decreased with 10 mg/kg of oxycodone (**Figure 6E**). If muscle rigidity begins at 3 mg/kg in females, perhaps the hyperlocomotion effect of oxycodone begins to be outweighed by the muscle rigidity effect at doses above 3 mg/kg. In other words, oxycodone may dose-dependently increase the drive to be mobile (hence the increase in open field distance covered at 10 mg/kg [10]) but by 10 mg/kg the muscle rigidity effect is strong enough to slow down the walking speed, so that the mice are not walking faster, they are simply walking for longer.

Ideally there would be a dose of oxycodone that would provide analgesia in incised animals without inducing tiptoe walking. Indeed, the effective dose of oxycodone can vary for different phenotypes of a single injury model; for example, in a mouse model of burn pain, the oxycodone dose required to reverse injury-induced CatWalk phenotypes was higher than the dose required to reverse injury-induced von Frey effects [39]. However, in the paw incision mice tested here, the oxycodone doses that were low enough not to affect paw print area in uninjured paws were not sufficient to reverse the paw print area phenotype in injured hind paws. The dose range at which analgesia is typically provided by oxycodone (1-20 mg/kg, depending on the injury model and sensory test [2; 14; 25; 29; 38; 39]) overlaps with the dose range for tiptoe walking shown here (1-10 mg/kg; **Figure 3**). Therefore, even though paw print area is a robust metric for demonstrating gait alterations induced by paw incision, this metric cannot be used to evaluate oxycodone’s ability to provide analgesia in this injury model.

Additional assessments are needed to confirm that the incision-induced paw print area reductions are indeed a pain-related phenotype, perhaps using other analgesics to reverse this effect on paw usage. Opioid treatment was shown to reverse paw usage impairments (guarding behavior) caused by paw incision [40] and paw print area changes induced by other injury models [1; 40] in rats. Further, paw print area changes induced by neuropathic injury were reversed by gabapentin in mice [21]. This suggests that altered paw placement is indeed a pain-related symptom that can be reversed by analgesics in rodents.

As a future direction, other metrics of paw usage could be analyzed to determine if they are altered by paw incision surgery and/or opioid treatment in mice. Whereas the CatWalk system measures two-dimensional paw placement with reliable precision, the amount of weight placed on the paw can only be generally estimated based on the light intensity of the paw print. Oxycodone may not reverse the area of the injured paw that makes contact with the glass, but it may affect how much weight is being placed on that reduced foot print. Static or dynamic weight bearing assays may be used as an alternative method to find a paw usage phenotype that is altered in an injured state and reversed by oxycodone. Having a better understanding of how paw usage varies with injury and/or oxycodone treatment would also help determine if other paw-directed assays (e.g. hot plate) are indeed appropriate to use in these contexts.

These results have interesting implications for how behavioral effects of opioids are studied in mice, including analgesia, muscle rigidity, and locomotion. Because both paw incision injury and opioid treatment cause mice to walk on the distal parts of their feet, paw print area is not a viable metric for demonstrating reversal of incision-induced paw usage with opioids. Therefore, alternatives to hind paw-directed reflexive assays for testing opioid analgesia, particularly in a post-surgical pain model, are still needed, but analysis of paw placement during walking may be an interesting area to look for new approaches. Overall, this research will help refine approaches to evaluating opioids in preclinical research, particularly when applied to mouse pain models.

## ACKNOWLEDGEMENTS

We would like to acknowledge funding for this work from the National Institute of Neurological Disorders and Stroke (R01 NS042595; RWG), National Institute of General Medical Sciences (T32 GM108539; VEB), and Washington University in St Louis (Dr. Seymour and Rose T. Brown Professorship in Anesthesiology; RWG). We have no conflicts of interests to report.

